# Analysis of Amino Acid Variants in Malignant Melanoma Cells Resistant to BRAF inhibition

**DOI:** 10.1101/2020.12.15.422879

**Authors:** Marisa Schmitt, Tobias Sinnberg, Katrin Bratl, Katharina Zittlau, Claus Garbe, Boris Macek, Nicolas C. Nalpas

## Abstract

Analysis of patient-specific nucleotide variants is a cornerstone of personalised medicine. Although only 2% of the genomic sequence is protein-coding, mutations occurring in these regions have the potential to influence protein structure and may have severe impact on disease aetiology. Of special importance are variants that affect modifiable amino acid residues, as protein modifications involved in signal transduction networks cannot be analysed by genomics. Proteogenomics enables analysis of proteomes in context of patient- or tissue-specific non-synonymous nucleotide variants. Here, we developed a proteogenomics workflow and applied it to study resistance to serine/threonine-protein kinase B-raf (BRAF) inhibitor (BRAFi) vemurafenib in malignant melanoma cell line A375. This approach resulted in high identification and quantification of non-synonymous nucleotide variants and (phospho)proteins. We integrated multi-omic datasets to reconstruct the perturbed signalling networks associated with BRAFi resistance and to predict drug therapies with the potential to disrupt BRAFi resistance mechanism in A375 cells. Notably, we showed that aurora kinase A (AURKA) inhibition is effective and specific against BRAFi resistant A375 cells. Furthermore, we investigated nucleotide variants that interfere with protein post-translational modification (PTM) status and potentially influence cell signalling. Mass spectrometry (MS) measurements confirmed variant-driven PTM changes in 12 proteins; among them was the runt-related transcription factor 1 (RUNX1) displaying a variant on a known phosphorylation site S(Ph)276L. We confirmed the loss of phosphorylation site by MS and demonstrated the impact of this variant on RUNX1 interactome.

## Introduction

Accumulation of mutations is one of the hallmarks of cancer cells and malignant melanoma is a type of cancer with the highest frequency of somatic mutations (1). Recent investigations showed that mutations of key signalling pathways in malignant melanoma are associated with poor clinical outcome, for example in the mitogen-activated protein kinase (MAPK)/extracellular signal-regulated kinase (ERK) pathway (2). The RAS/BRAF/MEK/ERK pathway is mutated to an oncogenic form in 30% of all cancers, with non-synonymous BRAF mutations in up to 50% of cutaneous melanomas (3). The predominant BRAF mutation is within the kinase domain with a single nucleotide substitution of valine to glutamic acid at amino acid 600 (4). This mutation can result in a 500-fold increased, dimerization-independent activation of BRAF, and thus leads to a constitutive activation of downstream signalling in cancer cells (3, 5). Targeted inhibition of the mutated BRAF kinase with selective inhibitors like vemurafenib, dabrafenib or encorafenib (BRAFi) results in a reduction of MAPK pathway signalling (5). However, almost all patients rapidly develop resistance to BRAFi monotherapy after a period of approximately five months (2). The considerable majority of BRAF resistance development is caused by molecular or genetic alterations that lead to MAPK pathway reactivation. The identification of multiple cellular mechanisms of resistance has greatly improved the understanding of malignancy and clinical outcomes of BRAF^V600E^ metastatic melanoma e.g. by the introduction of combined BRAF and MEK inhibition. However, variants that alter the corresponding protein modification status and therefore influence resistance, remain largely elusive. In addition, the precise effect of single nucleotide variant (SNV), insertion, deletion and frameshift variants at the proteome and PTM level are still largely unknown.

The past decade has seen a revolution in high-throughput sequencing technologies, which provide information on DNA/RNA sequence, gene structure and expression (6). Mass spectrometry (MS)-based proteomics is experiencing a technological revolution similar to that of the high-throughput sequencing. The current state-of-the-art “shotgun” proteomics workflows are capable of routine, comprehensive analysis of proteomes (7, 8) and posttranslational modifications (PTMs) such as phosphorylation (9, 10). However, most of the standard proteomics approaches identify peptides and proteins by matching MS/MS spectra against protein databases derived from public repositories (e.g. UniProt) that are not “individualised”, i.e. do not contain sequence information specific for an individual patient, tissue or cell line. Commonly used protein databases therefore inherently prevent identification of individual non-synonymous variants. Proteogenomics addresses this issue by combining nucleotide and protein sequencing information; this enables simultaneous study and integration of DNA sequence, RNA expression and splicing, protein isoform abundance, as well as localisation of protein PTMs in personalised fashion (11, 12).

Here we applied a proteogenomics approach to a single immortalised human melanoma cell line, A375, in its parental as well as in BRAFi resistant state, to analyse non-synonymous variants and their impact on signal transduction networks in context of acquired resistance to kinase inhibitors. Integration of matching genomics and (phospho)proteomics datasets allowed the reconstruction of signal transduction networks specific to individual cell lines. Subsequently, we were able to prioritise a number of drugs based on their disruptive potential on the signal transduction networks associated with resistance to BRAFi. We further investigated the impact of non-synonymous amino acid variants on phosphorylation sites and their putative functional effects on cell signalling.

## Experimental procedures

Only star methods are presented below; the rest of the methods is described fully in Supplementary Information.

### Experimental design and statistical rationale

For the whole-exome sequencing (WES), DNA was extracted from A375 sensitive (S) and resistant (R) cells. Since WES was used only for variant calling, one replicate was analysed for each cell line. Total genomic DNA was enriched for exome regions and sequenced on Illumina HiSeq 2000. Obtained reads were aligned to *H. sapiens* reference genome (GRCh38) using the HiSAT2 aligner. Variants were called using GATK software and incorporated into cell line-specific protein sequence database using in-house script.

The (phospho)proteomics data is derived from two sets of samples prepared and analysed by liquid chromatography-tandem mass spectrometry (LC-MS/MS). For the first screen, a total of 114 runs were performed with 60 min gradient for fractionated proteome and phosphoproteome measurements on an Exploris mass spectrometer. A375 S and A375 R cells were used for proteome and phosphoproteome measurements (three biological replicates per cell line, with 19 samples per replicate consisting of nine fractionated proteome samples and ten rounds of phosphopeptide enrichment). For the second screen, a total of 21 runs were performed with 60 min gradient for immunoprecipitation and 90 min gradient for peptide pulldown measurements on Q Exactive HF-X and HF mass spectrometers. Immunoprecipitation assays of Flag-tagged RUNX1, SILAC labelled A375 S RUNX1_KO cells transfected with pCMV_Flag_RUNX1 plasmid, pCMV_Flag_RUNX1_S276L plasmid or with empty vector plasmid (pCMV_Flag) were used (‘light’: ‘medium’: ‘heavy’) (three biological replicates). For synthetic peptide pulldowns in A375 S cells, label-free quantification between three independent replicates was performed (nine samples). Beads only were used as a control (three replicates).

LC-MS/MS raw data were processed using MaxQuant software (version 1.6.8.0 and 1.5.2.8). Statistical analyses were performed with Perseus (version 1.6.0.5) for the (phospho)proteome datasets (t-test and fisher exact test).

Biological assays were performed in three biological and six technical replicates, so that appropriate statistical analysis could be performed. Statistical analysis was performed with two-tailed unpaired t-test in GraphPad Prism (version 8). Separate controls were included in each experiment.

### Cell culture

The human metastatic BRAF^V600E^-mutated melanoma cell line A375 (CRL-1619, ATCC) was used in this study and authenticated by Microsynth AG. The generation of the cell line with acquired resistance to vemurafenib analogue PLX4720 (Selleckchem) (for simplicity referred to as “vemurafenib” in the Results section) was conducted as described previously (13). A375 S and R cells were grown in RPMI medium (Sigma-Aldrich) supplemented with foetal bovine serum (FBS, 10%, PAN Biotech) and penicillin/streptavidin (100 U/ml, PAN Biotech) at 37°C and 5% CO2.

For immunoprecipitation assays, SILAC-labelling of cells was performed as described previously (14) and detailed description of labelling of cells, CRISPR/Cas9-mediated knockout of RUNX1 and interaction assays can be found in the **Supplementary Information**.

### Incorporation of non-synonymous variants into protein databases

To integrate the proteogenomics datasets, we used an in-house bioinformatics pipeline, which is coded entirely in the R programming language (15). The transcript nucleotide sequences were extracted from GRCh38 *H. sapiens* genome assembly and Ensembl transcript annotation (via BSgenome and GenomicFeatures packages). These sequences were then *in silico* translated (from start to first stop codon) into a reference protein sequences database (Biostrings package). The called variants, within Variant Call Format files from A375 R and A375 S, were injected into each overlapping reference transcript nucleotide sequences and then *in silico* translated. The resulting protein sequences were written into two FASTA files containing reference variant protein sequences and sample-specific alternate variant protein sequences.

### Annotation of the biological impact of detected variants

In the current study, we prioritised amino acid variants based on their impact in context of BRAFi resistance in melanoma. For this purpose, known variant sites in melanoma, as well as known variant sites in cancer, were obtained from CGDS (16). These were overlapped with A375 identified variants and classified as loss/gain of sites. A list of oncogenes and tumour suppressor genes was compiled from Cosmic, ONGene, Bushman lab and Uniprot (17–19), whereas a list of genes harbouring somatic mutation in BRAFi resistant tumour was retrieved after reanalysis of published study (20). A375 variants found on these genes were annotated as relevant in cancer and/or BRAFi resistance.

A second impact scoring strategy was also performed to investigate protein phosphorylation-based signal transduction networks in A375 melanoma cells. Each reference/alternate variant protein sequence was annotated based on whether phosphorylation sites (S/T/Y) were lost and/or gained (IRanges package). A list of known kinase motifs was retrieved from PhosphoNetworks (21) and these motifs were searched along the reference/alternate variant protein sequences. Located kinase motifs were overlapped with variants position to determine loss/gain of the motifs. Known human phosphorylation sites were retrieved from PhosphoSitePlus and Phospho.ELM databases (22, 23). The variants identified in our study, which overlapped with known phosphorylation sites, were annotated as loss/gain of known phosphorylation. In a similar fashion, known variant sites in melanoma were obtained from CGDS (16) and overlapped with the variants from A375 R and S. A list of oncogenes and tumour suppressor genes was compiled from Cosmic, ONGene, Bushman lab and Uniprot (17–19). Variants on these genes that were identified in A375 R and S were annotated as cancer-relevant. A Levenshtein similarity score was calculated between reference and alternate variant protein sequences, whereby alternate sequences with less than 90% similarity to their reference were flagged.

Each amino acid within variant protein sequences were attributed a “+1” score for every overlap with an impact annotation. A summed score was then calculated for each amino acid within alternate variant sequence, and the maximum summed score was reported for that variant protein isoform. Because the score depends on the number of impacts used during the annotation, we also computed a scaled maximum score (between 0 and 1), to allow comparison between processings. Following the computation of all impacts, each variant protein isoform is ranked to allow prioritisation for follow up studies.

### Extraction and digestion of proteins

Cells were harvested at 80% confluence with lysis buffer (6 M urea, 2 M thiourea, 60 mM Tris pH 8.0) complemented with protease (complete Mini EDTA-free tablets, Roche) and phosphatase inhibitors (5 mM glycerol-2-phosphate, 5 mM sodium fluoride, and 1 mM sodium orthovanadate) and 1% N-ocetylglucoside (NOG, Sigma-Aldrich) for 10 min on ice. DNA and RNA were removed from the cell lysate using benzonase (1 U/ml, Merck Millipore) for 10 min on room temperature (RT). Cell debris was cleared by centrifugation (2,800 xg, 10°C, 20 min). Proteins were precipitated from cell lysates using eight volumes of acetone (−20°C) and one volume of methanol and incubated overnight at −20°C. The resulting solution was centrifuged (2,800 xg, 10°C, 20 min) to form a cell pellet. The pellet was washed two times with 80% acetone (−21°C) and resuspended in lysis buffer without NOG. Protein concentration was measured using Bradford assay. Extracted proteins (2-3 mg) were reduced with 10 mM of dithiothreitol (DTT) for 1 h, alkylated with 55 mM iodoacetamide for an additional hour and digested with Lys-C (Lysyl Endopeptidase, Wako Chemicals) for 3 h at RT. After adding four volumes of 10 mM ammonium bicarbonate, proteins were digested with trypsin (Promega Corporation) overnight. To stop the digestion, 1% trifluoroacetic acid (TFA) was added. Detailed description of high pH reverse phase chromatography can be found in the **Supplementary Information.**

### Phosphopeptide enrichment

Enrichment of phosphorylated peptides was performed using TiO_2_ beads (Titansphere, 10 μm, GL Sciences). TiO2 beads were resuspended in DHB solution (80% ACN, 1% TFA, 3% 2,5-dihydroxybenzoic acid (DHB)) and incubated for 20 min. Digested peptides were purified using Sep-Pak C18 Cartridge (Waters). In brief, Sep-Pak C18 Cartridges were activated with methanol and washed two times with 2% ACN, 1% TFA. After loading the sample, the column was washed again with solvent A (0.1% TFA) and eluted with 80% ACN, 6% TFA. Purified peptides were added to the TiO2 beads (beads to protein ratio, 1:2) and incubated for 10 min for each enrichment round (ten enrichment rounds). Phosphopeptide-bound TiO_2_ beads were sequentially washed with 30% ACN, 1% TFA, followed by 50% ACN, 1% TFA and 80% ACN, 1% TFA. Peptides were eluted with 5% NH_4_OH into 20% TFA followed by 80% ACN in 1% FA. The eluate was reduced by vacuum centrifugation, pH was adjusted to < 2.7 with TFA and peptides were desalted on C18 StageTips prior LC-MS/MS measurements.

### Liquid chromatography - mass spectrometry

Peptides were measured on an EASY-nLC 1200 ultra◻high◻pressure system (Thermo Fisher Scientific) coupled to a quadrupole Orbitrap mass spectrometer (Q Exactive HF and HFX and Exploris 480, Thermo Fisher Scientific) via a nanoelectrospray ion source. About 1 μg of peptides was loaded on a 20◻cm analytical HPLC◻column (75 μm ID PicoTip fused silica emitter (New Objective); in◻house packed using ReproSil◻Pur C18◻AQ 1.9◻μm silica beads (Dr Maisch GmbH)). LC gradient was generated by solvent A (0.1% FA) and solvent B (80% ACN in 0.1% FA) and 200 nl/min. Column temperature was kept at 40°C. For screen 1, all samples were measured on an Exploris 480 mass spectrometer using 60 min gradient for fractionated proteome (optimised gradient for each fraction) and phosphoproteome samples. The mass spectrometer was operated in data◻dependent mode, collecting MS spectra in the Orbitrap mass analyzer (60,000 resolution, 300-1750 *m/z* range) with an automatic gain control (AGC) set to standard and a maximum ion injection time set to automatic. For higher-energy collisional dissociation (HCD), the 20 most intense peptides were selected and fragment with a normalized collision energy of 28. MS/MS spectra were recorded with a resolution of 30,000 (fill time set to automatic). Dynamic exclusion was turned on and set to 30 sec. An *m/z* inclusion list (tolerance 10 ppm) was used to increase peptide coverage for RUNX1 and AURKA within the proteome and phosphoproteome measurements.

For the second screen, analysis of RUNX1 overexpression interactome (measured on Q Exactive HF-X), full MS were acquired in the range of 300-1750 *m*/*z* at a resolution of 60,000 (fill time 20 ms, AGC target 3E6). Twelve most abundant precursor ions from a survey scan were selected for HCD fragmentation (fill time 110 ms) and MS/MS spectra were acquired at a resolution of 30,000 on the Orbitrap analyzer. Precursor dynamic exclusion was enabled with a duration of 20 s (AGC target 1E5). Synthetic peptide pulldowns were analysed on Q Exactive HF mass spectrometer, full MS were acquired in the range of 300-1650 *m*/*z* at a resolution of 60,000 (fill time 25 ms, AGC target 3E6). MS/MS scans were acquired for the top seven most abundant precursor ions with a resolution of 60,000 and a fill time of 110 ms (AGC target 1E5).

### Mass spectrometry data processing

The raw data files were processed with the MaxQuant software suite (version 1.6.8.0 and 1.5.2.8) (24). The Andromeda search engine searched MS/MS data against *H. sapiens* reference (99,354 entries) and cell line-specific alternate databases (A375 = 29,104 entries), as well as UniProt *H. sapiens* (95,943 entries) database and commonly observed contaminants. Carbamidomethylation of cysteine (C) was set as fixed modification and oxidation of methionine, phosphorylation at serine, threonine or tyrosine were defined as variable modifications. Trypsin/P was selected as a protease. No more than two missed cleavages were allowed. The MS tolerance was set at 4.5 ppm and MS/MS tolerance at 20 ppm for the analysis using HCD fragmentation method. The false discovery rate (FDR) for peptides and proteins was set to 1%. For label-free quantification, a minimum of one peptide was required. For quantification of proteins in the immunoprecipitation experiments, the amino acids (Lys4)/(Arg6) and (Lys8)/(Arg10) were defined as ‘medium’ and ‘heavy’ labels for the comparison of RUNX1 overexpressed cell lines. For all other parameters, the default settings were used.

### Proteogenomics integration

Our in-house proteogenomics bioinformatic pipeline was used to integrate WES and MS datasets, specifically to check which mutations were identified across omic datasets. Initially, the reference and alternate variant protein sequences were *in silico* digested according to laboratory condition; i.e. digestion with trypsin and up to two missed cleavages (cleaver package). The overlap of MS-identified peptides with *in silico* digested peptides led to classification into reference (non-mutated peptide that overlap the mutation position on the reference protein), alternate (mutated peptide that overlap the mutation position on the alternate protein) or unspecific (non-mutated peptide that does not overlap any mutated positions) variant peptides. On the basis of this peptide classification, we summarised the peptides identification per variant protein isoforms, allowing coverage characterisation into reference only, alternate only, reference and alternate or unspecific. We finally focused on PTM (as implemented in the MaxQuant processing), which here consists in phosphorylation sites. Reference and/or alternate variant peptides found phosphorylated were flagged as such, as well as those were the phosphorylation occurred directly on the variant sites (either on reference or alternate variant sequences). This coverage information is exported within MaxQuant style processing results (tab-separated file as output).

Subsequently, we reconstructed the network of protein-protein (using BioGRID database) and drug-target (using DrugBank database) interactions; i.e. signalling network of BRAFi resistant A375 and RUNX1 interactome network (25, 26). The specificities of the drugs, interacting with nodes from the generated network, were calculated based on all possible target reported in DrugBank database. Drugs were prioritised further by summing the number of interactions their targets have within the network. The generated networks were exported (using igraph and RCy3 packages) into Cytoscape for further customisation (27).

## Results

To study the impact of amino acids variants on signal transduction networks, we selected the widely studied A375 melanoma cell line that harbours the BRAF^V600E^ mutation. We generated two closely related A375 cell lines, drug-sensitive (A375 S) and drug-resistant against the BRAF inhibitor vemurafenib (A375 R), as described previously (13). The resulting cell lines were subjected to whole exome sequencing, as well as proteomics and phosphoproteomics evaluation (**Figure S1A**). These omics measurements were then integrated to reconstruct signalling networks that are disturbed in BRAFi-resistant A375 cells and to investigate the impact of variants altering protein PTM status.

### Variance in protein abundance discriminates between BRAFi-sensitive and -resistant A375 cells

Initially, we investigated the mutational landscape of A375 R and A375 S cells by high-throughput sequencing. The majority (95%) of non-synonymous nucleotide variants consisted of nucleotide substitutions, the rest comprising frameshifts, deletions and insertions (**Figure 1A** and **Table S1.1**). This trend was consistent across both A375 cell lines. As expected, only 46 variants (out of 10,986) were classified as somatic in A375 R in comparison to A375 S cells, the rest being germline variants. Notably, most variants were already reported in Cosmic and/or dbSNP databases (**Figure S1B**), allowing for functional effect annotation. The variants affected genes, and in turn the corresponding proteins, from every subcellular compartment, such as cytoplasm (1067 variants), nucleus (680), mitochondrial matrix (229), mitochondrial import complex (115) and mitochondrial outer membrane (40). Analysis of the reference to alternate nucleotide substitution revealed a high frequency (64% of all substitutions) of adenine to guanine (and vice versa), as well as cytosine to thymine (and vice versa) exchange (**Figure S1C**). Subsequently, these non-synonymous variants were incorporated into their corresponding protein sequences. This led to the generation of several thousand additional protein isoforms contained within a cell-specific alternate protein sequence database (**Figure 1B**). Despite this large increase in the number of protein isoforms, the database search space increased by only 2%, which should result in no or very minor increase in false positive identification during MS spectral search (28).

**Figure 1:**
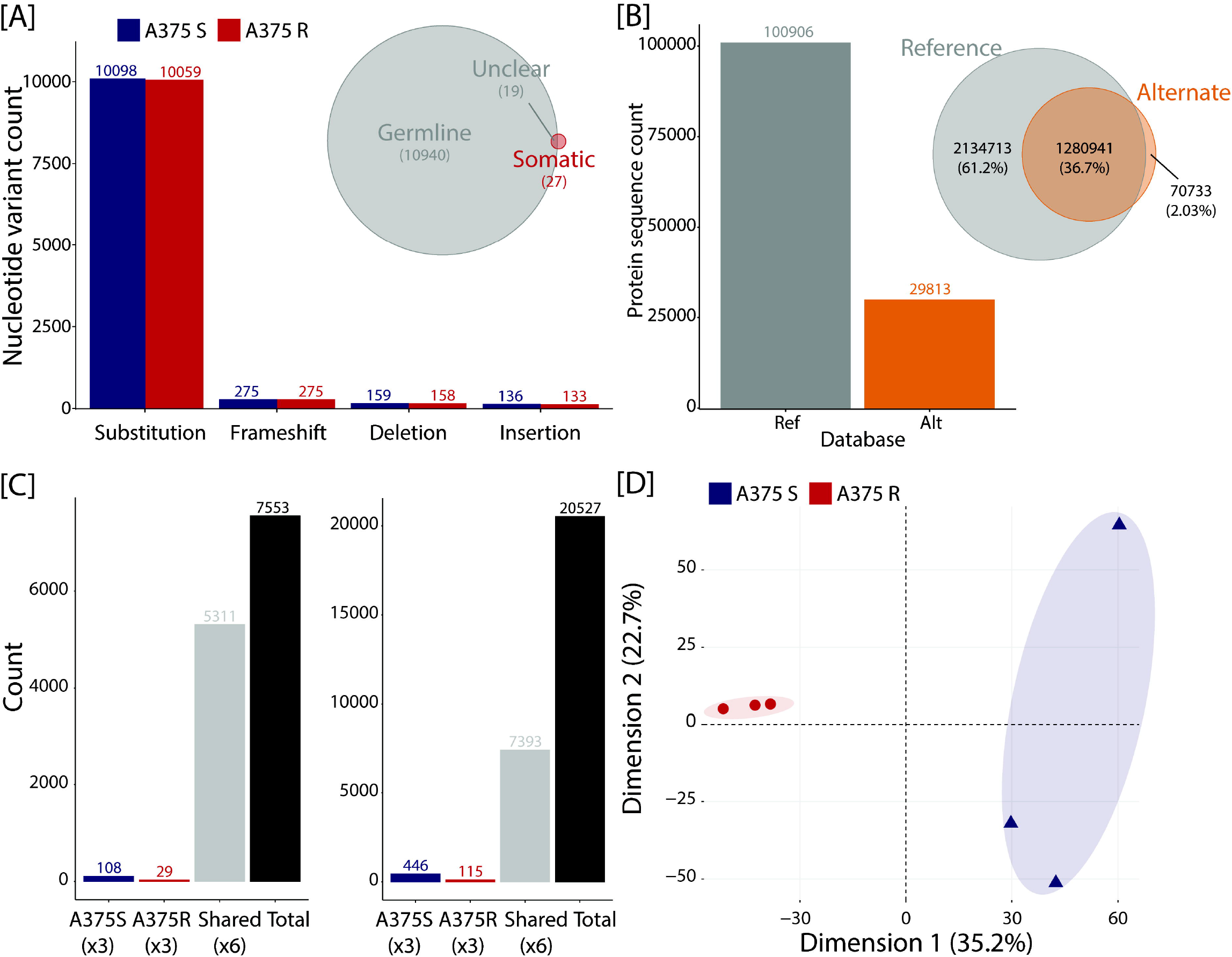
Multi-omics identification using proteogenomics. [A] Number of non-synonymous nucleotide variants is classified per type of alteration for A375 S and R cells. The venn diagram represents the overlap between germline and somatic non-synonymous nucleotide variants. [B] The number of protein sequences is displayed per reference (ENSEMBL) or alternate (WES from A375 R and S) databases, as well as the overlap in search space between databases (up to two missed cleavages). [C] Quantified protein groups and phosphorylated sites are counted based on whether they are shared or unique between A375 R and S cell lines. [D] Principal component analysis using protein abundances shows the separation of samples between cell lines (A375 R versus S), as well as the clustering of the biological replicates.

The alternate protein sequence database was then used, together with the reference database, for the processing of deep proteomics and phosphoproteomics data from A375 S and A375 R cells. High resolution MS led to the identification of more than 7,500 protein groups and 20,500 phosphorylated sites (**Figure 1C**), of which over 5,000 protein groups and 7,000 phosphorylated sites were quantified in all samples (n = 6). Using the quantitative data, these replicates were also characterised by high positive correlations at both the proteome (range 0.95 – 0.98) and phosphoproteome (range 0.73 – 0.90) levels (**Figure S1D**). Interestingly, a principal component analysis based on protein abundances revealed that the first dimension separated A375 R from A375 S replicates (**Figure 1D**). This separation was confirmed using the phosphorylated site abundances, although within the second dimension (**Figure S1E**). These initial quality controls highlighted the high measurement reproducibility for this dataset, as well as the ability to discriminate two closely related cell lines.

### Key signalling pathways are perturbed in BRAFi-sensitive and -resistant A375 cells

Subsequently, we investigated the non-synonymous nucleotide variants based on their functional effect as predicted by the oncoKB resource (29). This analysis revealed that five non-synonymous nucleotide variants resulted in a possible loss-of-function for cyclin dependent kinase inhibitor 2A (CDKN2A) and steroid receptor RNA activator 1 (SRA1), as well as a possible gain-of-function for aurora kinase A (AURKA) (**Figure 2A**). These variants were considered as potential driver mutations in context of resistance to BRAFi and retained for further analysis (30). To expand this driver list, we also overlapped the somatic non-synonymous nucleotide variants identified in A375 R against the study from Long and colleagues (20), who analysed 10 malignant melanoma patients (**Table S2.1**). While we did not observe any shared variants between A375 R and the patients’ data, we did find three shared genes harbouring different somatic variants (**Figure 2B**). Of note, the overlap of somatic mutations between patients in the Long *et al.* study was also very limited, which led us to retain all A375 R somatic variants as potential driver mutations for further analysis.

**Figure 2:**
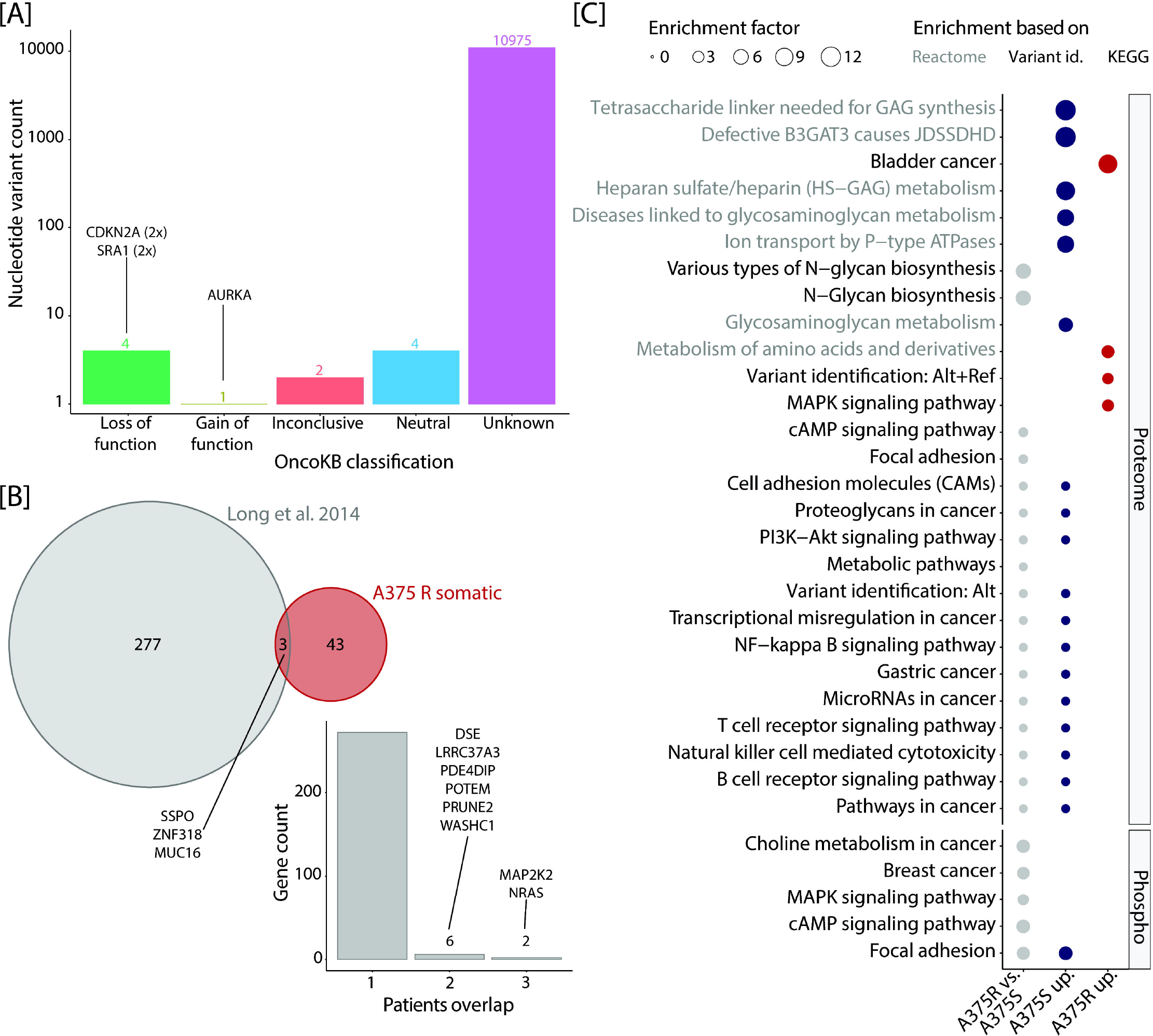
Comparison of BRAFi-sensitive and -resistant A375 cells at the genome and (phospho)proteome levels. [A] The non-synonymous variants are counted per functional effect category as predicted using the oncoKB resource (29). [B] A venn diagram represents the overlap in genes with non-synonymous somatic mutations between A375 R (identified in this study) and the patients from Long and colleagues’ study (20). Within the study from Long and colleagues, the genes with somatic non-synonymous nucleotide variants are represented as shared across one or more patients. [C] The signalling pathways (KEGG and Reactome) are displayed based on their overrepresentation using differentially changing proteins or phosphorylated sites (Fisher exact test FDR ≤ 0.2). The size of the pathways represents their enrichment factor. Pathway names are colour coded based on their functional database of origin (Reactome = grey, KEGG = black).

We next performed a quantitative comparison of A375 R versus A375 S using the protein and phosphorylated site abundances. MS analysis identified 359 significantly regulated proteins (*t*-test with s0 at 0.1 and FDR ≤ 0.05) between cell lines **(Figure S2A** and **Table S1.2)**, including the BRAF resistance marker protein nestin (NES), which was identified in our previous study (14). At the phosphoproteome level, we identified 187 differentially abundant phosphorylated sites (*t*-test with s0 at 0.1 and FDR ≤ 0.1) that were present on several key proteins, such as AP-1 transcription factor subunit (JUN) or eukaryotic translation initiation factor 4B (EIF4B) (**Figure** S**2B** and **Table S1.4**). Deeper functional characterisation was obtained through overrepresentation analyses based on different subset of significantly changing proteins and phosphorylated sites (Fisher exact test FDR ≤ 0.2). Notably the MAPK and cAMP signalling pathways, as well as focal adhesion, were overrepresented at both the proteome and phosphoproteome level (**Figure 2C**, **Table S1.3** and **S1.5**). We also identified several other pathways connected to cancer, immune response and glycoproteins (important for metastasis). Taken together, our data show that proteomics and phosphoproteomics provided meaningful insights into the disrupted signalling pathways that are implicated in BRAFi resistance.

### Amino acid variants are detectable within key signalling pathways and proteins

The aforementioned cell-specific protein database, used during the processing of the MS data, allowed us to detect a number of variant peptides, i.e. peptides harbouring reference or alternate amino acid. Among the MS-identified amino acid variants, most were shared across A375 R and S, a trend that was consistent at the proteome and phosphoproteome level (**Figure S2C**). Interestingly, these MS-detected variant protein isoforms were overrepresented in cancer-related, immune response and glycoprotein pathways (**Figure S2D** and **Table S1.6**). To confirm the quality of identification for these alternate variant peptides, we displayed the MaxQuant score, as well as intensity (mean and standard deviation) distribution, which was nearly identical to the rest of the peptides (**Figure S2E** to **F**). We also checked whether MS-identification of amino acid variants was dependent on variant genetic zygosity. Our data shows that the majority of alternate variant peptides were homozygous based on WES, however this trend did not translate to higher peptide intensities (**Figure S2G** to **H**).

Importantly, we could identify most of the commonly mutated genes in melanoma or in context of BRAFi resistance at the WES and/or (phospho)proteome levels (**Figure S2I** and **Table S1.7**). However, only the BRAF V600E variant could be identified at both the WES and proteome levels. Among the resistance marker genes found within the Long et al. study, only MAP2K3 (MEK3) harboured a variant in our WES dataset (R67W). These results highlight the capability of proteogenomics approach to detect amino acid variants on expressed proteins, which in turn reveal an overrepresentation of key cell signalling pathways.

### The perturbed signalling network in BRAFi resistant A375 cells can be targeted by several drugs

Subsequently, we integrated the significantly changing proteins and phosphorylation sites, as well as the list of potential driver mutations generated above, into a protein-protein interaction network (**Figure 3A, S3A** and **Table S1.8**). Only the top 200 entries were retained on the basis of their number of connections obtained from the BioGRID database. The size of the entries is scaled up using a custom bioinformatic script to represent their importance in context of cancer, melanoma and melanoma resistance (see **Supplementary Information**). A list of drugs was retrieved from DrugBank database and their targets were marked within this network (**Figure S3A**). Several entries were further highlighted through this approach due to their variant putative functional effect, such as AURKA (gain-of-function), CDKN2A and SRA1 (loss-of-function), CUL3, USP22 and MS4A1 (somatic mutation). Among these only AURKA and MS4A1 can be targeted by drugs, with AURKA displaying a relatively high betweenness centrality and number of degrees (**Figure 3A**).

**Figure 3:**
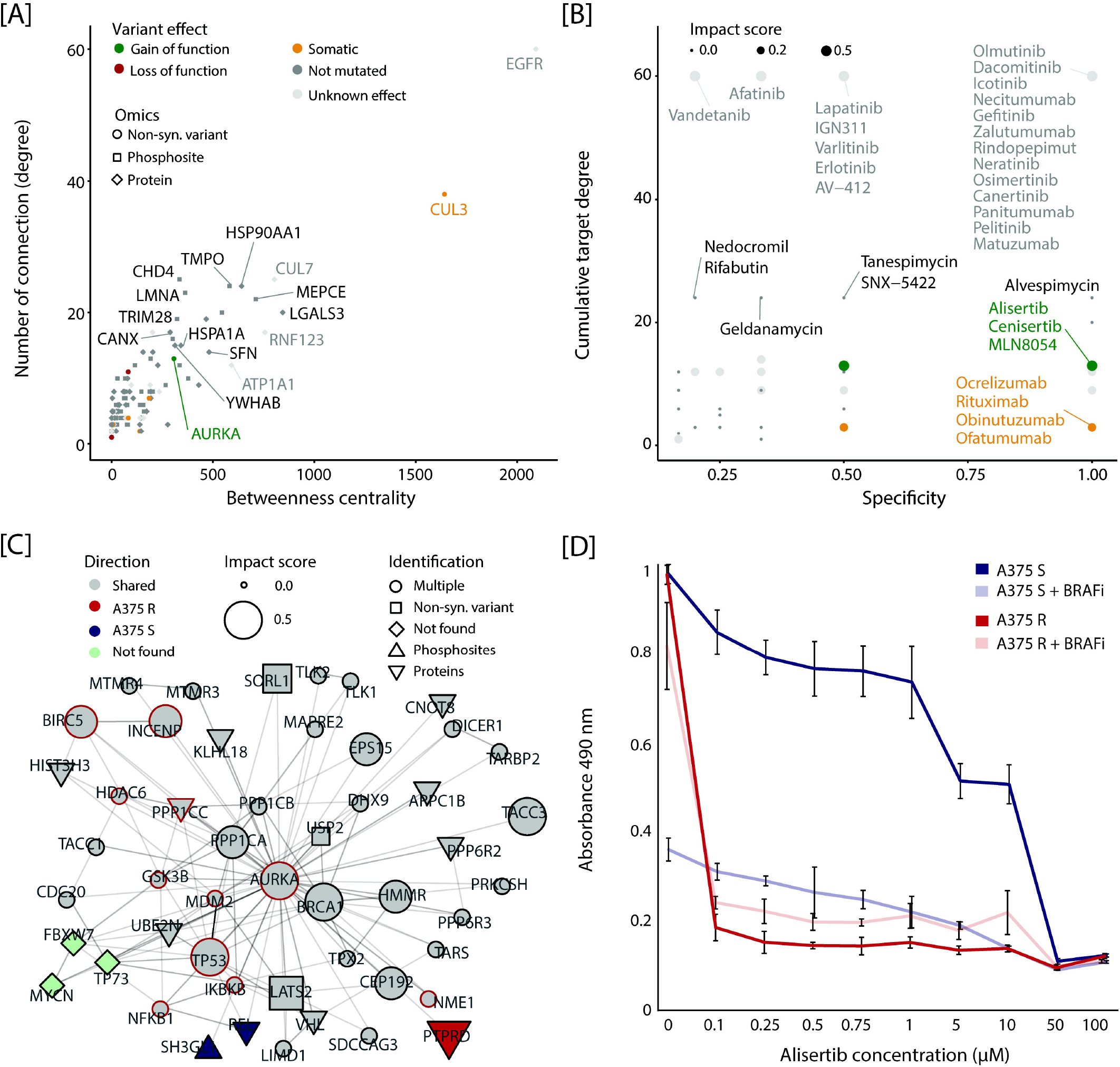
The disturbed signalling network in BRAFi resistant cells can be targeted by a number of drugs. [A] The interaction signalling network is generated based on list of putative driver mutations (circle), proteins (diamond) and phosphorylation sites (square). This schematic displays the distribution of nodes in function of their betweenness centrality and number of connections. Only the top 200 entries are displayed (ranked based on their interaction degree). Entries are coloured based on whether they harbour no variant (dark grey), a variant with unknown effect (light grey), a somatic variant (orange), a variant leading to loss-of-function (red) or a variant leading to gain- of-function (green). Entries that can be targeted by a drug are displayed with a red stroke. [B] The drugs, interacting with entries from the interaction signalling network, are displayed based on their specificity to their target and how many connections their targets have. Colour-coding corresponds to whether one of the drug targets harbour no variant (dark grey), a variant with unknown effect (light grey), a somatic variant (orange), a variant leading to loss-of-function (red) or a variant leading to gain-of-function (green). [C] The interaction signalling network allows visualisation of the top 50 interactors (ranked based on their interaction degree) of AURKA. The shape of the nodes specifies whether the node was identified harbouring a variant, quantified at the proteome level, quantified at the phosphoproteome level, found across more than one omics level or not found. Entries are coloured based on whether they were found significantly changing at the (phospho)proteome level, with increase abundance in A375 R (red) and A375 S (blue), not changing in abundance (grey) or not found (light green). Entries that can be targeted by a drug are displayed with a red stroke. The node size increases according to their importance in context of melanoma and BRAFi resistance (from 0 = no impact, up to 1 = high impact). [D] Cell viability assay of A375 S and A375 R cells treated with AURKA inhibitor alisertib at the indicated concentrations or in combination with the BRAF inhibitor vemurafenib (2 μM). Cell viability was determined with MTS assay 72h after treatment start (n = 6). Error bars represent standard deviations of replicates.

This allowed us to prioritise potential drugs based on their target specificity, as well as on the number of degrees their targets have with the rest of the signalling network (**Figure 3B**). Among the prioritised therapies were inhibitors of EGFR (e.g. olmutinib), HSP90AA1 (e.g. alvespimycin), AURKA (e.g. alisertib) and MS4A1 (e.g. rituximab). Because AURKA was the only variant annotated with a gain-of-function, as well as a relatively large signalling network (**Figure 3C**), we decided to experimentally validate the action of the compound alisertib on A375 R and S cells. We found that BRAFi-resistant A375 cells were sensitive to AURKA inhibition (AURKAi) with alisertib (31), regardless of the absence/presence of BRAFi vemurafenib (**Figure 3D** and **S3B**). Conversely, A375 S tolerated alisertib (in absence of BRAFi) and were able to proliferate. Our results demonstrate that the integration of proteogenomics datasets has the potential to predict effective and specific drug therapy in context of BRAFi resistance.

### Multiple amino acid variants directly affect protein phosphorylation status

We then investigated the elusive impact of amino acid variants on the phosphorylation status within A375 cell signalling. All non-synonymous variants were annotated based on whether they had an impact on S/T/Y amino acids, kinase motifs, known variants, known PTMs, known oncogenes and protein sequence changes superior to 90% of the reference protein (**Figure S4A** to **B** and **Table S1.9**). Interestingly, 6,103 variants resulted in a loss of S/T/Y residues, whereas 5,876 resulted in a gain, which represented a large potential for disrupting the phosphorylation-mediated cell signalling. Through investigation of the phosphoproteome dataset, we could confirm that 12 amino acid variants resulted in an actual loss or gain of phosphorylation events within A375 cells. To prioritise these variants, we annotated them using the PolyPhen software (as well as SIFT), which predicted a variant on RUNX1 protein (S276L) as likely damaging (**Figure 4A**). Notably, this variant site on RUNX1 was one of the few variant sites detected with a high localisation probability (**Figure 4B**). To further evaluate the quality of MS-identification, we compared the phosphorylated site intensities between the variant versus all other sites, which revealed lower intensity among variant sites (**Figure S4C**). Similarly, when displaying the MaxQuant score and localisation probability, only a few phosphorylated variant sites (including the site on RUNX1) could be considered as high confidence sites (**Figure S4D**). While these phosphorylated variant sites were not significantly changing in abundance between A375 R and S and thus are unlikely to be connected with BRAFi resistance, we hypothesised that they may still have a functional effect on RUNX1 activity and related cell signalling.

**Figure 4:**
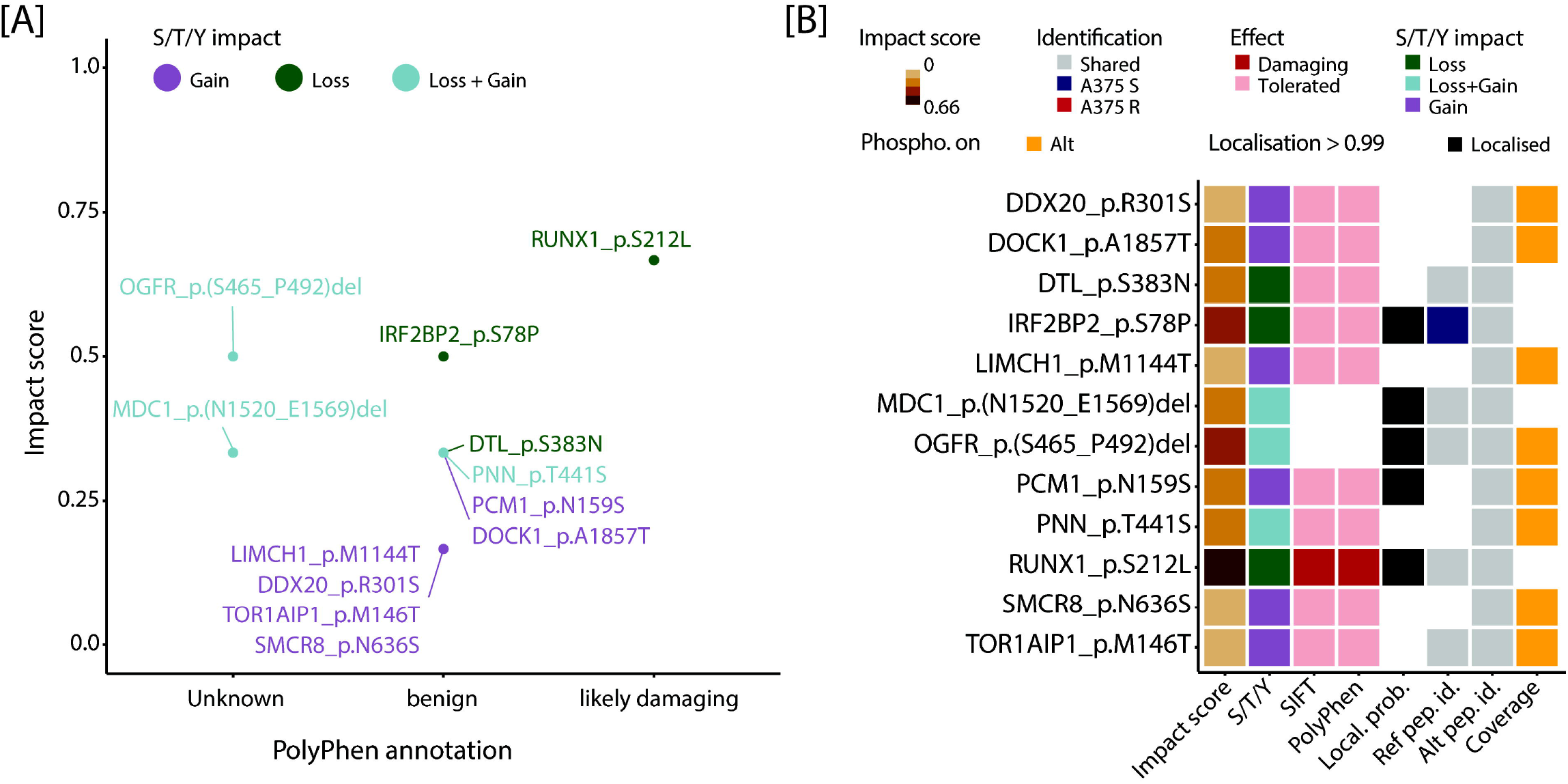
Multiple amino acid variants directly affect protein phosphorylation status. [A] Scatter plot of the non-synonymous amino acid variants that have an impact on protein phosphorylation status either as a loss of a S/T/Y amino acids (green), gain of S/T/Y (purple) or loss/gain of S/T/Y (blue). The PolyPhen annotation classifies these variants based on their possible effect on protein function, while the impact score prioritises these variants due to their importance in context of melanoma and phosphorylation status disruption. [B] Heatmap of the non-synonymous amino acid variants that have an impact of protein phosphorylation status. For each non-synonymous amino acid variant is displayed its impact factor in context of melanoma and phosphorylation status, the actual phosphorylation effect (i.e. loss, gain), the SIFT and PolyPhen annotation, whether the modification was localised (i.e. localisation probability ≥ 0.99), whether the reference and alternate variant peptide were identified exclusively in A375 R or A375 S or in both, and whether the modification is found on the alternate variant peptide (as opposed to the reference).

### Loss of a phosphorylation site on RUNX1 alters its interactome and transcriptional activity

We then focused on the variant impacting the phosphorylation status of RUNX1 protein, a key transcription factor involved in cell proliferation, differentiation and apoptosis (32). This amino acid variant led to the loss of a known phosphorylation site due to change from serine to leucine at position 276 (**Figure 5A**). The reference and alternate variant peptides were identified with high resolution MS in both A375 S and R cells (**Figure S5A** to **B**). We hypothesised that this variant is likely to influence the interactome of RUNX1. Therefore, we generated a RUNX1 gene knockout in A375 S cells using the CRISPR/Cas9 system. Single cell clones were selected for further analysis based on their effective RUNX1 knockout (KO). As a control we used a non-targeting (NonTar) control guide sequence. A375 RUNX1 KO cells showed an insertion of 215 bp in the Exon 1 of the gene compared to reference DNA of A375 S cells (**Figure S5C**). The lack of expression of RUNX1 protein was confirmed by western blot and MS analysis (**Figure S5D**). To study the impact of the loss of a modifiable amino acid, we performed immunoprecipitation of Flag-tagged RUNX1_wt and RUNX1_S276L in RUNX1 KO SILAC labelled cells in three independent replicates (**Figure 5B**, **Table S3.1** to **3.2**). The interactome analysis by LC-MS/MS revealed that RUNX1 and its core binding factor CBFB were significantly enriched in both pulldowns compared to Flag-empty vector (**Figure S5E** and **F**). Interestingly, the known interaction partner histone deacetyltransferase HDAC1 was enriched in RUNX1_wt interactome and depleted in the RUNX1_S276L interactome (**Figure 5B**).

**Figure 5:**
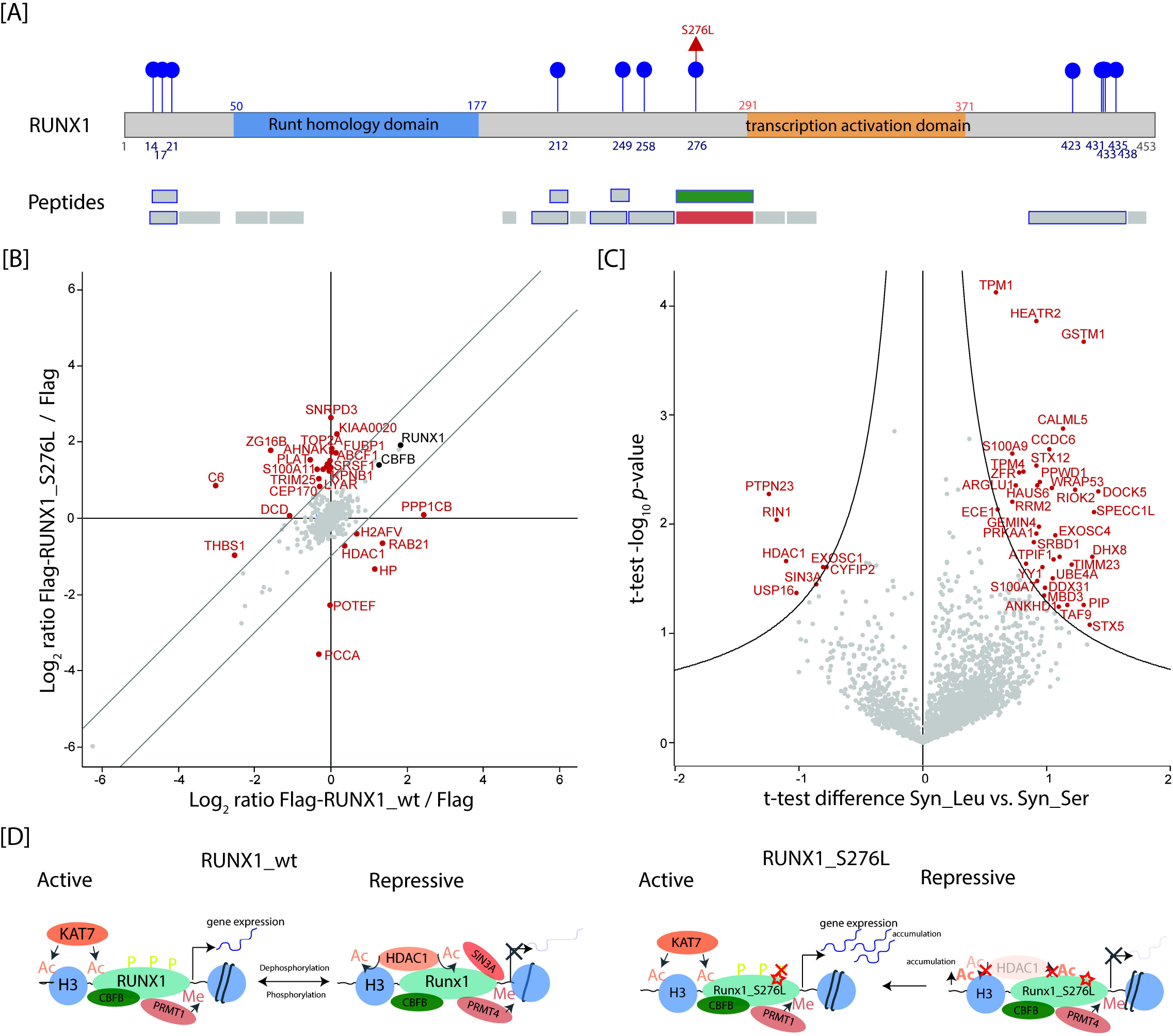
Loss of a known phosphorylation site leads to a change in RUNX1 interactome. [A] Schematic overview of the transcription factors RUNX1 protein. Numbers indicate the positions of amino acids residues within the protein. Identified phosphorylation sites are highlighted in blue and identified amino acid variants are highlighted in red. Identified peptides by LC-MS/MS are shown in the second panel. Phosphorylated peptides are indicated with a blue border, while reference and alternate variant peptides are highlighted in green and red, respectively. [B] Interaction proteomics screen in A375 RUNX1_KO cells stably overexpressing Flag-tagged RUNX1_wt or Flag-tagged RUNX1_S276L. SILAC protein expression (log_2_) of RUNX1_wt or Flag-tagged RUNX1_S276L relative to the corresponding control cell line (Flag tag only). RUNX1 and its core binding factor CBFB are marked in black. Significantly up and downregulated proteins are highlighted in red. Results represent three replicates per experiment group. [C] Volcano plot of synthetic alternate peptide (Syn_Leu) versus synthetic reference peptide (Syn_Ser) pulldowns of A375 cells. The log_2_ fold change in abundance between Syn_Leu and Syn_Ser are plotted against −log_10_ p-value (n = 3). Black lines indicate the significance threshold based on student t-test (FDR < 0.01; s0 = 1.2). Significantly up and downregulated proteins are highlighted in red. [D] Schematic overview of proposed interaction of RUNX1_wt and RUNX1_S276L with main transcriptional regulators.

To assess the impact of this variant on the interactome, we performed pulldown assays with synthetic peptides harbouring the amino acid sequence for reference and alternate variant of RUNX1 in A375 cells (**Figure 5C**, **Table S3.3** to **3.4**). As in the interactome study, HDAC1 was significantly depleted in the pulldown of alternate versus reference variant peptide indicating that the interaction between HDAC1 and RUNX1 is disturbed due to the variant. We also identified transcriptional repressor SIN3A to be significantly depleted in alternate variant peptide pulldown compared to reference pulldown similar to HDAC1. RIN1 and PTPN23 showed the same trend as HDAC1 and SIN3A; and both proteins are known to act as regulator of RAS-mediated mitogenic activity (33, 34). The proteins enriched in alternate pulldown compared to reference variant peptide pulldown were overrepresented in TGFβ signalling, melanogenesis and insulin signalling pathways (**Figure S5G** and **Table S3.5**). Taken together, we demonstrate that the loss of the known phosphorylation site on S276 has an impact on the interactome of RUNX1 (**Figure S5H**) and postulate that it leads to altered transcriptional activity of the protein (**Figure 5D**).

## Discussion

In this study, we used a common cell line model of melanoma (A375), a well-known cancer for its high mutation load (1) and the potential for rewiring cellular networks (12, 35). Two consortia, namely the Clinical Proteomic Tumor Analysis Consortium and The Cancer Genome Atlas, have greatly contributed to the development of onco-proteogenomics (11, 36, 37). However, proteogenomics studies are still relatively rare and, due to their complexity, out of reach of most proteomics (or genomics) laboratories. Here, we use such a proteogenomics workflow to analyse a single melanoma cell line in context of BRAFi resistance in order to predict tumour-specific drug therapies and to investigate variant impact on phosphorylation-mediated cell signalling.

### Proteogenomics reconstructs the signalling network linked to BRAFi resistance in A375 cells

In this study, the nucleotide variant analysis revealed very similar numbers across A375 cell lines, the large majority being SNVs, which is consistent with a previous study (38). We also observed characteristic nucleotide substitutions, whereby two thirds of substitutions are comprised of transitions. The C to T transition was highly represented and is known to result from sun-light exposure, which is highly relevant for skin cancer (39). The proteome coverage we obtained for A375 cells is similar to other state-of-the-art MS-based study of cancer cell lines (40). The differentially changing proteins and phosphorylated sites between A375 R and S revealed that the MAPK signalling, cAMP signalling and focal adhesion pathways were found overrepresented and are of critical importance for melanoma resistance to BRAFi (41). We then reconstructed the signalling network associated with BRAFi-resistance in A375 cells; i.e. using the putative driver mutations, as well as the differentially abundant proteins and phosphorylation sites. This approach highlighted several hubs and high impact entries, such as (1) variants leading to gain-of-function on AURKA, loss-of-function on CDKN2A and SRA1 and somatic mutations on CUL3, USP22 and MS4A1; (2) increase in protein abundance within A375 R for EGFR and increased within A375 S for HSP90AA1; and (3) increase in phosphorylated sites abundance within A375 S for JUN. Among these entries, several are known for their involvement in melanoma susceptibility, development, resistance and therapy (42–48). Several drugs were identified and ranked based on their potential to disrupt this signalling network; notably alisertib, a highly specific inhibitor of AURKA, which has been previously reported for its beneficial effect in combination with BRAF and MEK inhibitors in melanoma treatment (31, 42). We experimentally validated the use of alisertib on A375 S and R cells and could show that A375 R cells are sensitive to AURKAi. Our data confirm that AURKA has a critical role in the context of resistance and may be suitable for the treatment of melanoma as reported previously (49).

### Proteogenomics pinpoints several peptides that are phosphorylated on the variant site

We investigated further the amino acid variants that could be confirmed by MS and may be important in melanoma development or resistance. Over 500 protein groups and 158 phosphorylated sites were found with at least one alternate variant peptide. Our identification results are in the same range (or higher) as other studies investigating amino acid variants using custom protein sequence databases (50, 51). Interestingly, an overrepresentation analysis of alternate variant proteins revealed cancer-related, immune response and glycoprotein pathways (52–56). There are two possible explanations for such result, first it suggests an accumulation of variants on proteins belonging to these pathways, as these variants would provide a survival advantage for cancerous cells, and second the proteins harbouring these variants are highly abundant in A375 cells, thus facilitating amino acid variant detection (30, 51).

As observed in our dataset, many variants may have an impact on the post-translational modification of proteins. Around 14.8% of all amino acids in the human proteome are serine, threonine or tyrosine (19), which are predominantly modified by phosphorylation. Several studies have reported that these three amino acids are disproportionally affected by missense mutations (57, 58). While these may not all be relevant in tumour cells, since not all genes are expressed at any one-time, previous studies have shown the deleterious effect of such variants (11, 12). Here, we confirmed by MS a change in phosphorylation status for 12 variant protein isoforms, either as a loss or gain of phosphorylated sites. Several of these protein isoforms were key cell signalling molecules and may be involve in cancer development, for example MDC1 (59), OGFR (60) and PCM1 (61). Interestingly, we also confirmed by MS the presence of a variant on RUNX1, which was annotated as likely damaging due to its known role in cancer and loss of phosphorylation site at position S276 (62). The identification of these variants by MS is certainly very encouraging and further work is needed to increase the identification score and localisation probability of the alternate variant peptides, as well as to investigate their function.

### Rewiring of signal transduction network due to loss of a known phosphorylation site on RUNX1

We experimentally validated this striking example of a loss of a known phosphorylation sites on RUNX1 and showed that this variant has an impact on the interactome of RUNX1. The transcription factor RUNX1 is mutated in 3.03% of melanoma patients and so far, 43 mutations are described in the literature for cutaneous melanoma (63). The variant site S276L of RUNX1 is located in a highly modified region of the protein and may influence the nearby transcriptional activation domain. Wee *et al.* showed *in vitro* that the triple phosphorylation at the sites S249, T273 and S276 are important for the interaction with the histone acetyltransferase p300 and thus lead to the regulation of gene transcription via chromatin remodelling (62). Here, we could not identify p300 in the interactome studies of RUNX1 by immunoprecipitation of overexpressed RUNX1 or synthetic peptide pulldowns. However, we identified the transcriptional activator WWTR1 (TAZ) and KAT7 and the corresponding transcriptional repressors HDAC1 and SIN3A to be changing between reference and alternate pulldown of RUNX1. The loss of the interaction to HDAC1 by mutating RUNX1 at S48, S303 and S424 to aspartic acid *in vitro* has been described previously (64). Here, we hypothesised that the interaction is associated with the modification status of the protein. The crosstalk between acetylation/deacetylation mediated and phosphorylation/dephosphorylation may alter the transcriptional activity by RUNX1. It is well known in the literature that RUNX1_wt switches between active and repressive state due to modifications like acetylation and phosphorylation and binding of interaction partners like HDAC1, PRMT1 and P300 (65). We postulate that RUNX1_S276L remains in the active state due the loss of binding to transcriptional repressors like HDAC1 and SIN3A, which could lead to the accumulation of acetylation on the protein itself as well as histones. This may result in stronger transcriptional activity, which should be tested in further experiments. Taken together, we propose that this variant changed the interactome of RUNX1 and altered the transcriptional activity of RUNX1.

## Conclusions

Proteogenomics is a powerful tool to study the mode of action of disease-associated mutations at the genome, transcriptome, proteome and PTM level. Here, we applied a proteogenomics workflow to study the melanoma cell line A375 sensitive and resistant to BRAF inhibition. The investigation and integration of multi-omic datasets allowed us to reconstruct the perturbed signalling networks associated with BRAFi resistance. This resulted in the prioritisation of key actionable nodes and the prediction of drug therapies with the potential to disrupt BRAFi resistance mechanism in A375 cells. Notably, we demonstrated the use of AURKA inhibitor as an effective and specific drug against BRAFi resistant A375 cells. We also detected the loss or gain of several phosphorylation events due to variants. We could confirm the loss of Ser276 phosphorylation site by MS as a direct consequence of variant S276L on the transcription factor RUNX1. Our results suggest that this mutation has an impact on the interactome of RUNX1 and may be responsible for change in its transcriptional activity. We believe that such proteogenomics workflow is readily applicable to other types of cancer and within cell lines, as well as patient-derived samples.

## Abbreviations

A375 R: BRAFi-resistant A375 cells
A375 S: BRAFi-sensitive A375 cells
AGC: automatic gain control
AURKA: Aurora kinase A
AURKAi: AURKA inhibition
BRAF: Serine/threonine-protein kinase B-raf
BRAFi: BRAF inhibitor
DHB: dihydroxybenzoic acid
ERK: Extracellular signal-regulated kinase
FBS: Foetal bovine serum
FDR: false discovery rate
HCD: higher-energy collisional dissociation
LC-MS/MS: liquid chromatography-tandem mass spectrometry
MAPK: Mitogen-activated protein kinase
MS: Mass spectrometry
NOG: N-ocetylglucoside
PTM: Post-translational modification
RT: Room temperature
RUNX1: Runt-related transcription factor 1
SNV: Single nucleotide variant
WES: Whole-exome sequencing

## Data availability

The high throughput nucleotide sequencing data have been deposited in the NCBI Sequence Read Archive (66) with the bioproject accession number PRJNA616103. The mass spectrometry proteomics data have been deposited to the ProteomeXchange Consortium via the PRIDE (67) partner repository with the dataset identifier PXD018305. The WES bioinformatics pipeline is available online (68).

## Supplemental data

This article contains supplemental data.

## Author contributions

M.S., T.S., B.M. and N.C.N. designed the study. M.S. and K.Z. performed the proteomics experiments, while K.B. helped with the interaction proteomic screen. M.S. and N.C.N. analysed the data and performed statistical analysis. M.S. and N.C.N. wrote the manuscript with the input from all authors.

## Acknowledgements

The authors acknowledge Prof. Dr. Yulia Skokowa for fruitful discussions, Prof. Dr. Stefan Stefanovic for synthetic peptides, Prof. Dr. Stefano Stifani for the pCMV_RUNX1 expression plasmid, Dr. Karsten Krug, Lena Thiess and Payal Nashier for their help and advice during the project, as well as c.ATG Core Facility in Tuebingen for the WES library preparation and sequencing. This work was supported by the High Performance and Cloud Computing Group at the Center for Data Processing of the University of Tuebingen, the state of Baden-Wuerttemberg (bwHPC), Deutsches Konsortium für Translationale Krebsforschung (DKTK), German Research Foundation (DFG) grants No. INST 37/935-1 and INST 37/741-1 FUGG (to B.M.), and by intramural funding from the University of Tuebingen for the promotion of junior researchers (to N.C.N).

