## Supplementary information for "Analysis of Amino Acid Variants in Malignant Melanoma Cells Resistant to BRAF inhibition"

### Supplementary Materials and Methods

#### Cell culture

For immunoprecipitation assays of Flag-tagged RUNX1, cells were grown in RPMI-1640 SILAC (Sigma-Aldrich) lacking arginine, lysine and leucine. Leucine (12.5 mg/ml, Sigma-Aldrich), penicillin/streptomycin (100 U/ml, PAN), dialyzed FBS (10% (v/v), PAN) and stable isotope-encoded arginine and lysine were added to the SILAC medium. The ‘light’ SILAC media was further supplemented with L-[12C6,14N2] lysine (Lys0) and L-[12C6,14N4] arginine (Arg0) (Cambridge Isotope Laboratories), whereas L-[2H4] lysine (Lys4) and L-[13C6] arginine (Arg6) were added to the ‘medium’ SILAC media and L-[13C6,15N2] lysine (Lys8) and L-[13C6,15N4] arginine (Arg10) to ‘heavy’ SILAC media.

#### DNA extraction and Whole Exome Sequencing

Cells were harvested by centrifugation (80×g, 3 min) and washed twice with PBS. DNA was extracted using QIAamp DNA Mini (QIAGEN) according to the manufacturers’ instructions. WES libraries were prepared using SureSelect Human All Exon V5 Rapid Run (Agilent) according to manufacturers’ instructions. Paired-end sequencing was performed on an HiSeq 2500 instrument (Illumina) with on average >80 reads per base. The WES measurements were performed at c.ATG Core Facility in Tuebingen. Raw sequence data were then processed using an in-house pipeline developed at the Proteome Center Tuebingen according to GATK best guidelines (1).

#### WES data analysis

Raw sequence data were processed using an in-house pipeline developed at the Proteome Center Tuebingen. The raw reads were initially quality checked using FastQC software (2). Illumina adapters and 5’/3’ low quality bases were trimmed from reads using Trimmomatic (3). Paired-end reads from individual libraries were then aligned to *H. sapiens* reference genome (GRCh38) using the HiSAT2 aligner (4). Reads resulting from PCR duplication were marked using Picard package. Variants were called using the GATK HaplotypeCaller (germline) and Mutect2 (somatic) workflow (1). Variants were recalibrated for score and filtered (soft-filter) using GATK. OncoKB software was used to perform the annotation and functional effect prediction of detected variants (5).

Reanalysis of the WES data from Long and colleagues (6) was performed using the same bioinformatics pipeline described above.

#### Generation of RUNX1 knockout melanoma cell line using CRISPR/Cas9

*RUNX1* gene knockout was carried out by CRISPR/Cas9-mediated genome editing according to the published protocol (7). The SpCas9 plasmid PX459 (Plasmid 62988) was obtained from a non-profit plasmid share repository (Addgene). Suitable CRISPR target sites within *RUNX1* Exon 1 positive strand were identified using the ‘CRISPR Design Tool’ (<http://crispr.mit.edu/>). The respective oligonucleotide inserts (ITD) (5’- CACCGGATGAGCGAGGCGTTGCCGC *-*3*’* (forward), 5’- AAACGCGGCAACGCCTCGCTCATCC *-*3*’* (reverse)) were designed with overhangs compatible for ligation into PX459 linearised by digestion with BbsI (New England BioLabs). Oligonucleotides were phosphorylated with polynucleotide kinase T4 PNK (New England BioLabs), annealed and inserted into the plasmid using T4 DNA ligase (New England BioLabs) and transformed into chemocompetent DH5α E. coli cells (New England Biolabs). Oligonucleotide inserts (ITD) (*5’-gtattactgatattggtggg-3’* (forward), *5’-cccaccaatatcagtaatac-3’* (reverse)) were designed as CRISPR/Cas9 non-targeting (NonTar) control sgRNA and cloned into the SpCas9 plasmid PX459. Melanoma cells were seeded with low density (100,000 cells per ml), grown for 24h in RPMI-1640 medium without FBS. Transfection of the SpCas9/sgRNA plasmid or SpCas9/NonTar plasmid was carried out with Lipofectamine 2000 (Thermo Fischer Scientific) according to the manufacturer's instructions. On the next day, cells were selected using puromycin (2 µg/ml, Invivogen) and incubated for two days. Once individual colonies formed, single colonies were picked, cultured in separate wells and expanded in 6-well plates until cell number was sufficient for further analysis. Genomic DNA was isolated using GeneElute Mammalian Genomic DNA MiniPrep Kit (Sigma-Aldrich) and PCR amplification was performed using primers (5′- GGCCAGTACCTTGAAAGCGA -3′ (forward) and 5′- TGGTAGGAGCTGTTTGCAGG -3′ (reverse)) and sequenced by Sanger sequencing.

#### Western blot

Cells were harvested in lysis buffer and proteins were precipitated overnight with acetone/methanol (-20°C). Protein extracts were separated on 4–12% NuPAGE Bis-Tris gels (Novex, Life Technologies), transferred to PVDF membranes (0.2 µm, Sigma-Aldrich). The blot membranes were blocked in 1% (v/v) Tween-20 and probed with primary antibody followed by horseradish peroxidase-conjugated secondary antibodies. Primary antibodies used were anti-RUNX1 (PA519638, Thermo Fisher Scientific) and anti-Histone H3 antibody (D1H2) (#4499, Cell Signaling Technologies). Secondary antibodies used were anti-rabbit IgG, HRP-conjugated (#7074, Cell Signaling Technologies) and anti-mouse IgG, HRP-conjugated (#7075, Cell Signaling Technologies). ECL was detected by exposure with the Fusion SL instrument (Vilber Lourmat).

#### Overexpression and immunoprecipitation of RUNX1

Immunoprecipitation (IP) of overexpressed Flag-tagged RUNX1 in A375 S RUNX1_KO SILAC cells was performed with Flag M2 antibody (F3165, Sigma-Aldrich) in three biological replicates. Site-directed mutagenesis, in order to generate pCMV_Flag_RUNX1_S276L, was performed using QuickChange II Site-Directed Mutagenesis Kit (Agilent Technolgies), according to the manufacturer’s instructions. For control experiments, pCMV_Flag lacking RUNX1 mRNA sequence was used. A375 S RUNX1_KO SILAC cells were transfected with pCMV_Flag_RUNX1 plasmid, pCMV_Flag_RUNX1_S276L plasmid or with empty vector plasmid (pCMV_Flag) using Lipofectamine 2000 transfection reagent (Thermo Fisher Scientific). For IPs of RUNX1, 1 mg of each SILAC labelled cell line were mixed according to the protein amount. Cell lysates were precleared for 1h at 4°C with washed Pierce Protein G magnetic beads (Thermo Fisher Scientific) using 5 µl per mg lysate. Flag M2 antibody was coupled to the beads by incubation at 4°C for 20 min in incubation buffer (50 mM Tris-HCl pH 8.0, 300 mM NaCl, 1 mM EDTA, 0.5% (v/v) Triton X100). The beads were washed three times with DPBS to remove unbound peptides. Precleared cell lysates and Flag-tagged Pierce Protein G magnetic beads were incubated for 2h at 4°C while shaking in incubation buffer supplemented with protease inhibitor (complete Mini EDTA-free tablets, Roche) and phosphatase inhibitor buffers (5 mM glycerol-2-phosphate, 5 mM sodium fluoride, and 1 mM sodium orthovanadate). Pierce Protein G magnetic beads were used as control. Beads were washed three times with incubation buffer and two times with DPBS. Proteins were eluted by incubation at 95°C for 10 min in NuPAGE LDS sample buffer (4x) (Thermo Fisher Scientific).

Protein samples from immunoprecipitation experiments were prepared for LC-MS/MS using in gel digestion. Proteins were separated on a NuPAGE Bis-Tris 4-12% gradient gel (Thermo Fisher Scientific) and stained with Coomassie Brilliant Blue solution. Gel lanes were cut into small pieces and washed three times with washing buffer (5 mM AmBiC, 50% (v/v) ACN) to remove Coomassie stain. To reduce disulfide bonds, 10 mM dithiothreitol (DTT) in 20 mM AmBiC was added and incubated at 56°C for 1h. After alkylation of the disulfide bonds with iodoacetamide (55 mM (IAA) in 20 mM AmBiC) for 30 min at RT, gel pieces were washed with washing buffer and dehydrated with 100% (v/v) ACN and vacuum centrifugation (10 min). Proteins were digested with trypsin (12.5 ng/ml in 20 mM AmBiC, Promega Corporation) at 37° overnight. Digested peptides were extracted from gel pieces with 3% (v/v) TFA in 30% (v/v) ACN, followed by 0.5% (v/v) acetic acid in 80% (v/v) ACN and 100% (v/v) ACN. All extracts were combined, concentrated by vacuum centrifugation and purified on C18 StageTips.

#### Pulldown assays with synthetic peptides and on-bead digestion

Synthetic peptides comprising 17 amino acids and a biotinylated linker in the N-terminus were dissolved in DPBS and 750 µg of each peptide was used for three independent pulldowns. Peptides were coupled to Pierce streptavidin magnetic beads (100 µg peptides/ 1 mg beads) (Thermo Fisher Scientific) by incubating beads with an excess of synthetic peptides for 2h at RT in pulldown buffer (50 mM Tris-HCl pH 8.0, 300 mM NaCl, 1 mM EDTA, 0.5% (v/v) Triton X-100). The beads were washed four times with DPBS to remove unbound peptides. Cell extracts of A375 S were pre-cleared by incubation at 4°C for 30 min with washed Pierce streptavidin magnetic beads using 5 µl per mg lysate. For each peptide pulldown 2 mg of pre-cleared input material and beads without synthetic peptides were used as a control. Cell extracts and peptides were incubated for 1h at RT while shaking in incubation buffer supplemented with protease inhibitor and phosphatase inhibitor buffers (5 mM glycerol-2-phosphate, 5 mM sodium fluoride, and 1 mM sodium orthovanadate). Beads were washed four times with DPBS and two times with H_2_O. Bound proteins from synthetic peptide pulldowns were digested directly on the beads. Denaturation buffer (8 M urea, 2 M thiourea, 50 mM Tris-HCl pH 8.0) and 1 mM DTT was added to beads and incubated for 1h at RT while shaking. After adding of 5.5 mM of IAA and an incubation step of 1h at RT, proteins were pre-digested with Lys-C (1 µg, Lysyl Endopeptidase, Wako Chemicals) for 3h at RT. Samples were diluted four times with 20 mM AmBiC and digested with trypsin (1 µg, Promega Corporation) overnight at RT. Digested proteins in the supernatant were transferred to a new tube, acidified using 10% (v/v) TFA and purified on C18 StageTips.

**Table 1: List of used synthetic peptides in this study.**

| **Peptide** | **Sequence** |
| --- | --- |
| RUNX1_S276 | SGSGSPSVHPATPISPGRASGM |
| RUNX1_L276 | SGSGSPSVHPATPILPGRASGM |

Kind gift of Prof. Dr. Stefan Stefanovic, University of Tuebingen

#### Synchronisation of cells and cell viability assay

To synchronize cells in G1/S boundary, cells were seeded at 30-40% confluency in RPMI-1640 medium and exposed to two sequential thymidine block (8). Briefly, cells were exposed to 2 mM thymidine for 18 h, incubated in fresh medium for 9 h and again exposed to 2 mM thymidine (Sigma-Aldrich) for 18 h. The block was released by adding fresh medium to cells.

For cell viability assay, cells were seeded (2 x 10^3^) into 96-well plates. After four hours, media was changed and cells were incubated for 72 h with increasing concentrations of Alisertib (0.1 µM to 100 µM, MLN8237, Selleckchem) or with DMSO. In addition, cells were incubated in presence or absence of the BRAF inhibitor vemurafenib (2 µM, PLX4720, Selleckchem). After washing with DPBS, cell viability was assessed by CellTiter 96^®^ Aqueous One Solution Cell Proliferation Assay (Promega Corporation), according to the manufacturer’s instructions. Briefly, 10 µl of the MTS reagent was added to the cells in culture medium and incubated for 1 h at standard culture conditions. The absorbance was read at 490 nm with Infinite M Nano microplate reader (Tecan group Ltd.).

#### High pH reversed-phase chromatography

Peptides were fractionated using Pierce high pH reversed-phase peptide fractionation Kit (Thermo Fisher Scientific), according to the manufacturer’s instructions. Briefly, 100 µg of peptides were fractionated into nine fractions based on the hydrophobicity using a step gradient generated with ACN (5%-50% in 10 mM NH_4_OH). The pH of the fractions was adjusted to <2.7 with TFA, dried by vacuum centrifugation and purified on C18 StageTips prior LC-MS/MS measurements.

#### Significance testing and pathway analysis

Statistical analyses were performed with Perseus software suite (version 1.6.5.0). For the (phospho)proteome investigation of BRAFi resistance in A375 cells, the drug-sensitive (n = 3) and drug-resistant (n = 3) A375 cells were compared using (1) label-free quantification for the proteins and (2) intensities for the phosphorylation sites. Each omics dataset was analysed separately and entries were filtered out if not quantified in all samples. Additionally, the reverse and potential contaminants were filtered out from the protein and phosphorylation site datasets. Notably, the phosphorylation sites intensities were normalised by the corresponding proteins intensities. A t-test was used to compute p-values and identify significantly changing entries between A375 R and S samples. The p-value were corrected for multiple testing with a permutation-based FDR (s0 = 0.1 and FDR ≤ 0.05 [proteome] or FDR ≤ 0.1 [phosphoproteome]). The proteins and phosphorylation sites, which were statistically tested in Perseus, are listed in **Table S1**.

For proteomic interaction studies of RUNX1, protein groups were kept for further statistical analysis only if quantified in 3 out of 3 replicates. The SILAC ratios of the three independent replicates were averaged and an arbitrary cut-off of two-fold change was used to determine significant SILAC ratios. The log_2_ transformed ratios were plotted against intensities (log_10_). For synthetic peptide pulldowns, label-free quantification between three independent replicates was performed and ratios were subjected to t-test analysis, with a permutation-based FDR threshold of 0.01 and s0 value of 1.2. A list of known interaction partners of RUNX1 was retrieved from BioGrid and mapped to the dataset. A list of all protein identifications is provided in **Table S3**.

The resources used for annotation of proteins were Kyoto Encyclopaedia of Genes and Genomes (KEGG), Gene Ontology Biological Function (GOBP) and Reactome Pathway database (Reactome). The fisher exact test (FDR ≤ 0.2 [BRAFi resistance studies] or FDR ≤ 0.1 [interaction studies]) was used to test for overrepresented functions or pathways among significantly changing entries against the background of identified entries. The displayed pathways were selected based on highest FDR or enrichment score. A list of all overrepresentation results is provided in **Table S1 and S3**.

### Supplementary figures

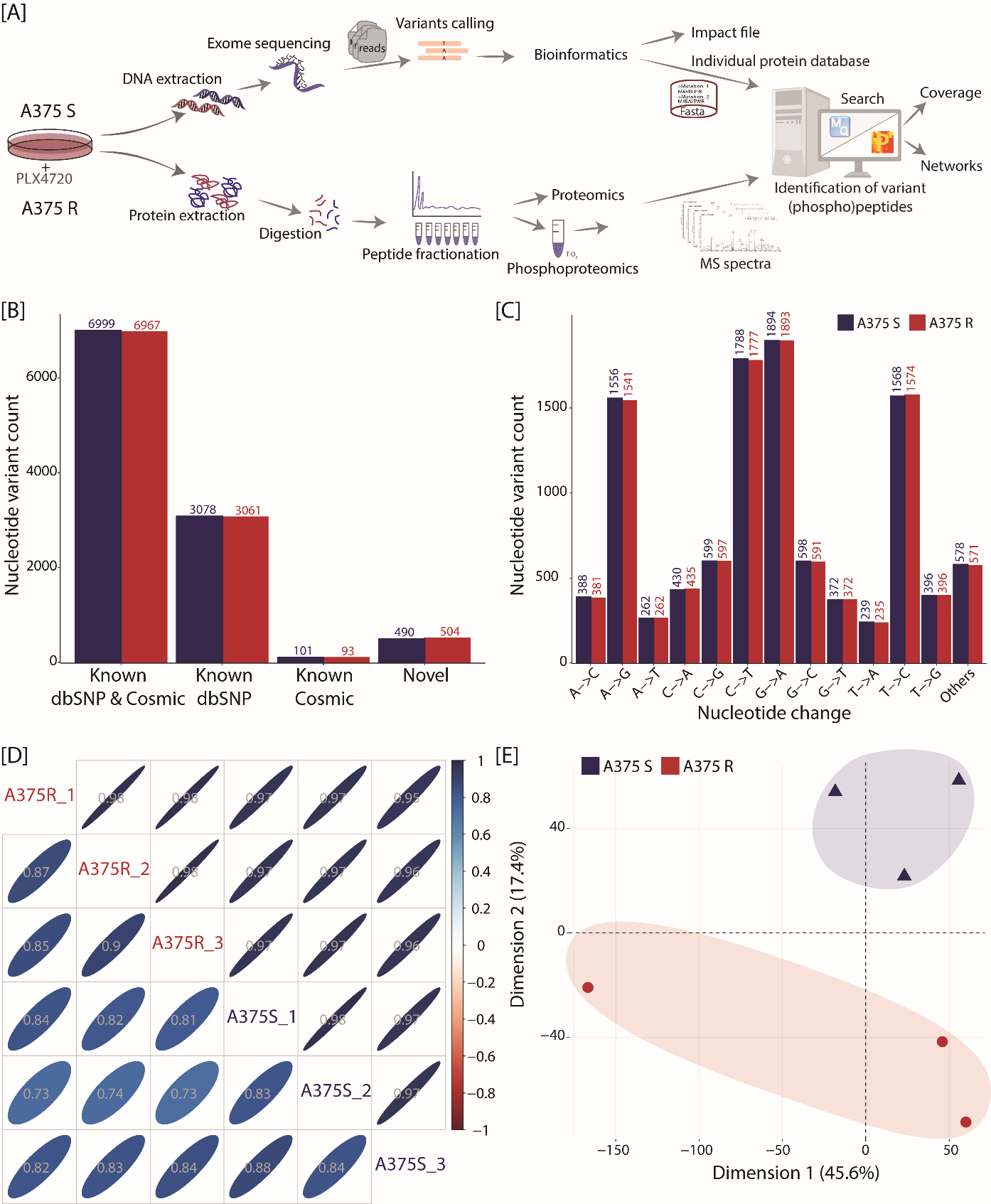

#### Figure S1: Multi-omics identification using proteogenomics.

[A] Schematic overview of the proteogenomics workflow. Vemurafenib (PLX4720) sensitive and resistant melanoma cell line A375 were used in this study. For the WES, DNA was extracted and sequenced on Illumina HiSeq 2000. Variants were called using GATK software and incorporated into cell line-specific protein sequence database using in-house script. For the proteomics and phosphoproteomics workflow, cells were lysed and proteins were digested using trypsin. The resulting peptide mixture was fractionated using Pierce high pH reversed-phase peptide fractionation Kit. Fractions were measured directly (proteome) or applied to phosphopeptide enrichment using titanium dioxide (TiO_2_) prior to LC-MS/MS. MS raw data were processed with MaxQuant software and further analysed using in-house bioinformatic pipeline and Perseus. [B] Exome sequencing results of A375 S and R stratified based on whether these are reported in COSMIC or dbSNP databases or are novel variants. [C] Count of non-synonymous nucleotide variants transitions and transversions for A375 S and R. [D] The correlation between A375 R and S samples (n = 3 each) is represented based on protein (above diagonal) and phosphorylated sites (below diagonal). The correlation score corresponds to Spearman's rank correlation coefficient. [E] Principal component analysis using phosphorylated site abundances shows the separation of samples between cell lines (A375 R versus S).

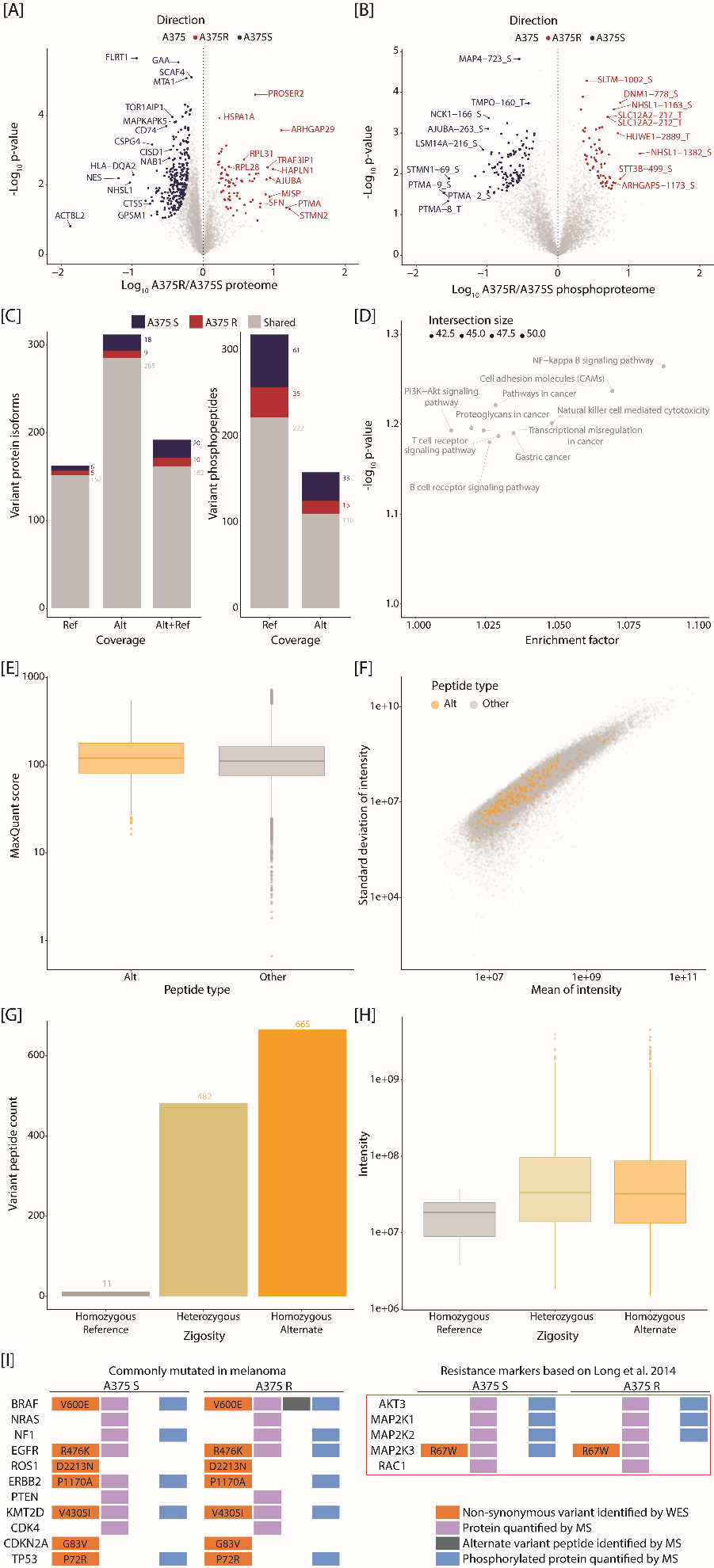

#### Figure S2: Comparison of BRAFi-sensitive and -resistant A375 cells at the genome and (phospho)proteome levels.

[A and B] Comparison of A375 R vs. A375 S based on quantitative proteome [A] and phosphoproteome [B] data. Significantly changing proteins [A] or phosphorylated sites [B] are coloured in red and blue when increasing in A375 R and A375 S, respectively (*t*-test with s0 at 0.1 and FDR ≤ 0.05 [proteome] or ≤ 0.1 [phosphoproteome]). [C] The identification of reference and alternate variant isoforms is displayed at the proteome and phosphoproteome levels. Variant protein isoforms or variant phosphorylated sites were either quantified in both phenotypes (grey) or exclusively detected in A375 R (red) or A375 S (blue). [D] The overrepresented KEGG pathways is based on alternate variant protein isoforms identified in either A375 R or S. [E] The distribution of MaxQuant score is displayed per alternate variant (orange) or all other (grey) peptides. [F] The distribution of peptide average intensity is plotted against the corresponding intensity standard deviation. The colour coding differentiates alternate variant (orange) versus all other (grey) peptides. [G] The identified alternate variant peptides are counted based on their genetic zygosity (as determined by WES). [H] The distribution of alternate variant peptide intensities is displayed per genetic zygosity (as determined by WES). [I] The most common known driver mutations, as well as resistance markers (based on Long *et al.* 2014), for malignant melanoma are selected. Each variant identification status is displayed based on WES and LC-MS/MS.

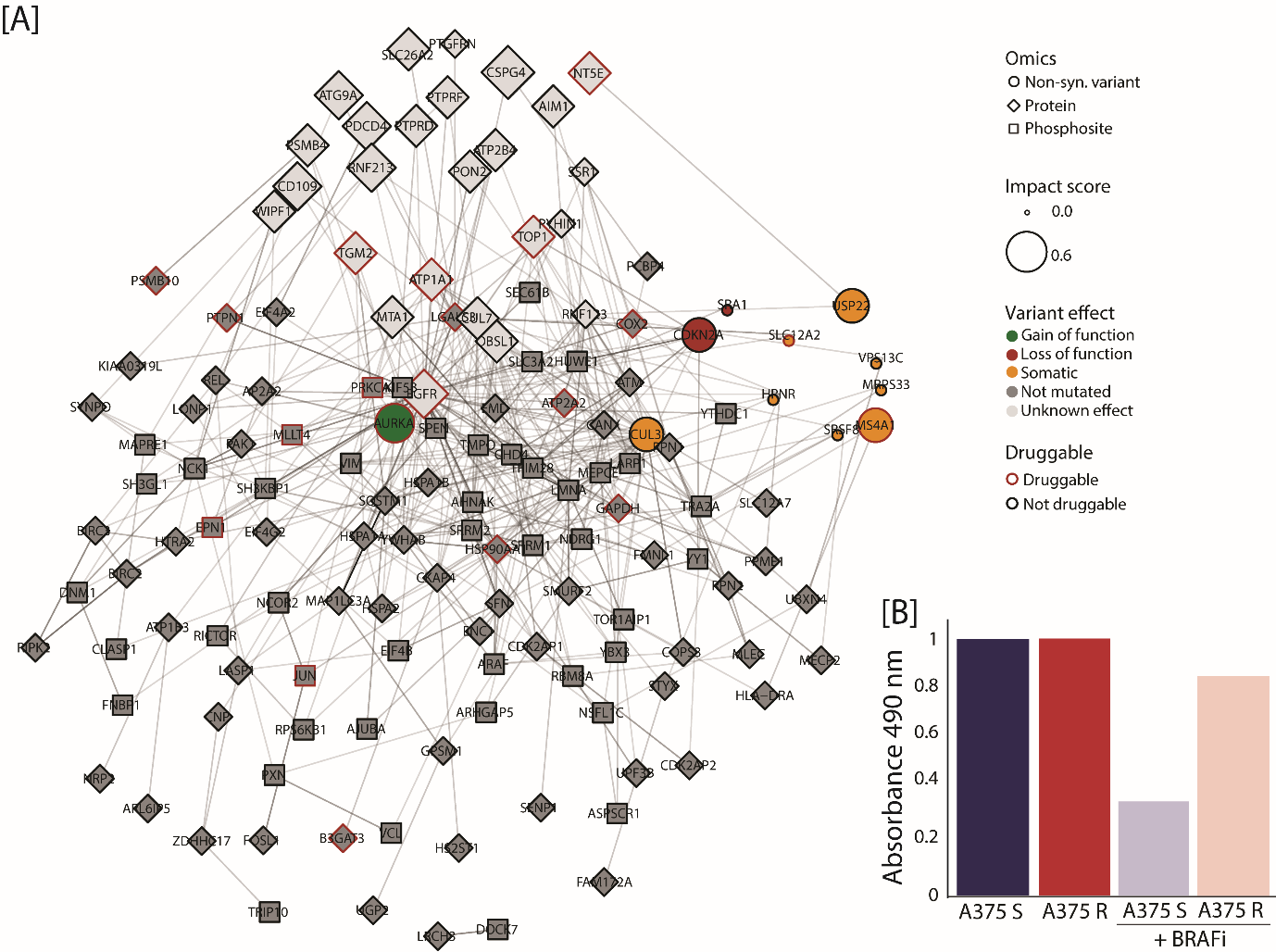

#### Figure S3: The disturbed signalling network in BRAFi-resistant cells can be targeted by a number of drugs.

[A] The interaction signalling network is generated based on list of putative driver mutations (circle), proteins (diamond) and phosphorylation sites (square). Only the top 200 entries are displayed (ranked based on their interaction degree). Entries are coloured based on whether they harbour no variant (dark grey), a variant with unknown effect (light grey), a somatic variant (orange), a variant leading to loss-of-function (red) or a variant leading to gain-of-function (green). Entries that can be targeted by a drug are displayed with a red stroke. Entries were prioritised based on their importance in context of melanoma and BRAFi resistance (from 0 = no impact, up to 1 = high impact) and their node size increased accordingly. [B] Cell viability assay of A375 S and A375 R cells treated with AURKA inhibitor Alisertib at the indicated concentrations or in combination with the BRAF inhibitor vemurafenib (2 µM). Cell viability was determined with MTS assay 72h after treatment start (n = 6).

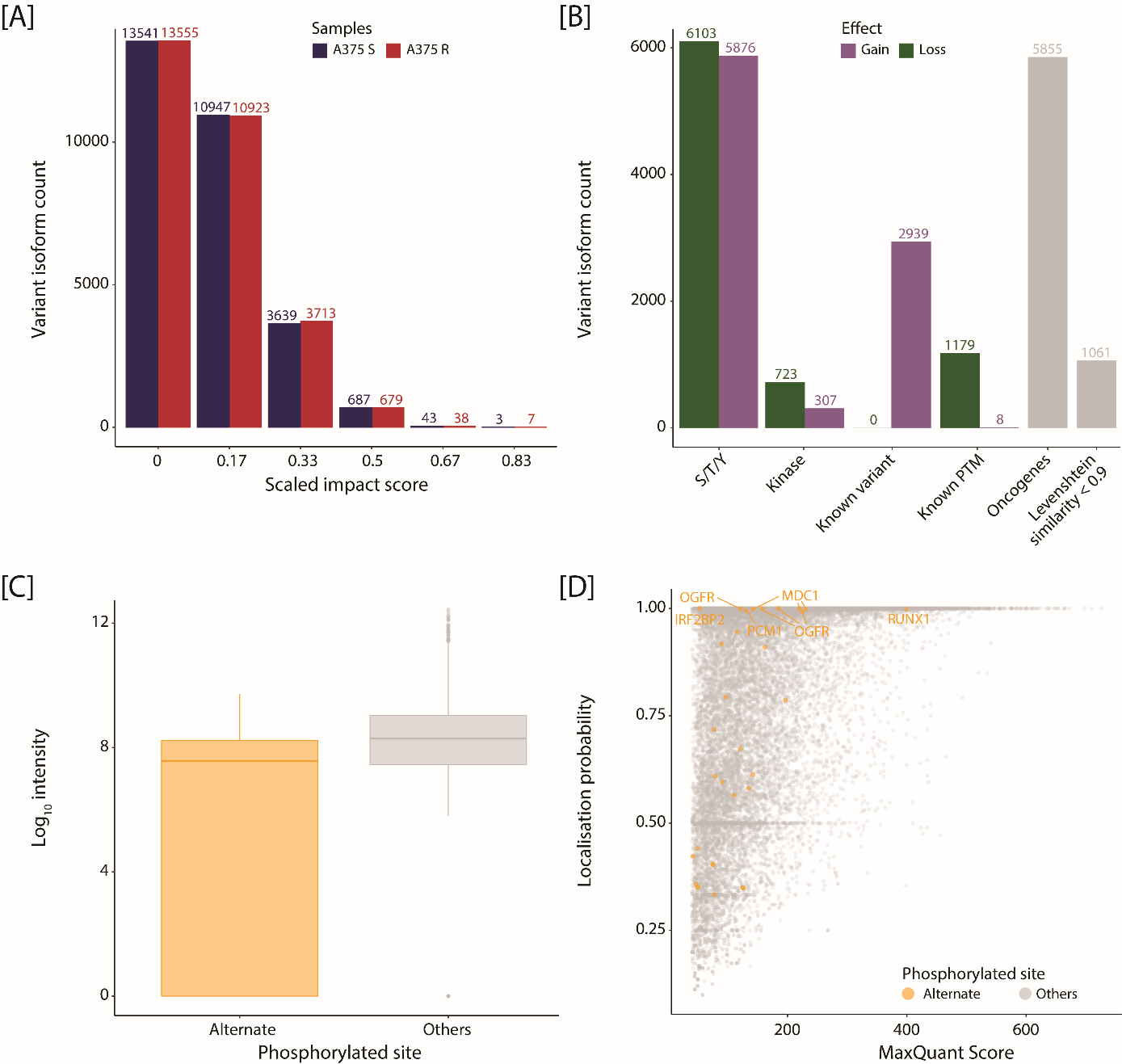

#### Figure S4: Multiple amino acid variants directly affect protein phosphorylation status

[A] The variant protein isoforms are counted per impact score representing their importance in context of melanoma and phosphorylation status (from 0 = no impact, up to 1 = high impact). The count is performed separately for A375 R (red) and S (blue). [B] The variant protein isoforms are counted for each individual component entering in the computation of the impact score. These components comprise impact on S/T/Y amino acids, kinase motifs, known variants, known PTMs, known oncogenes and protein sequence changes superior to 90% of the reference protein. [C] The distribution of phosphorylation site intensities is plotted. The colour coding differentiates alternate variant (orange) versus all other (grey) phosphorylated sites. [D] The distribution of phosphorylated site MaxQuant score is plotted against the corresponding localisation probability. The colour coding differentiates alternate variant (orange) versus all other (grey) phosphorylated sites.

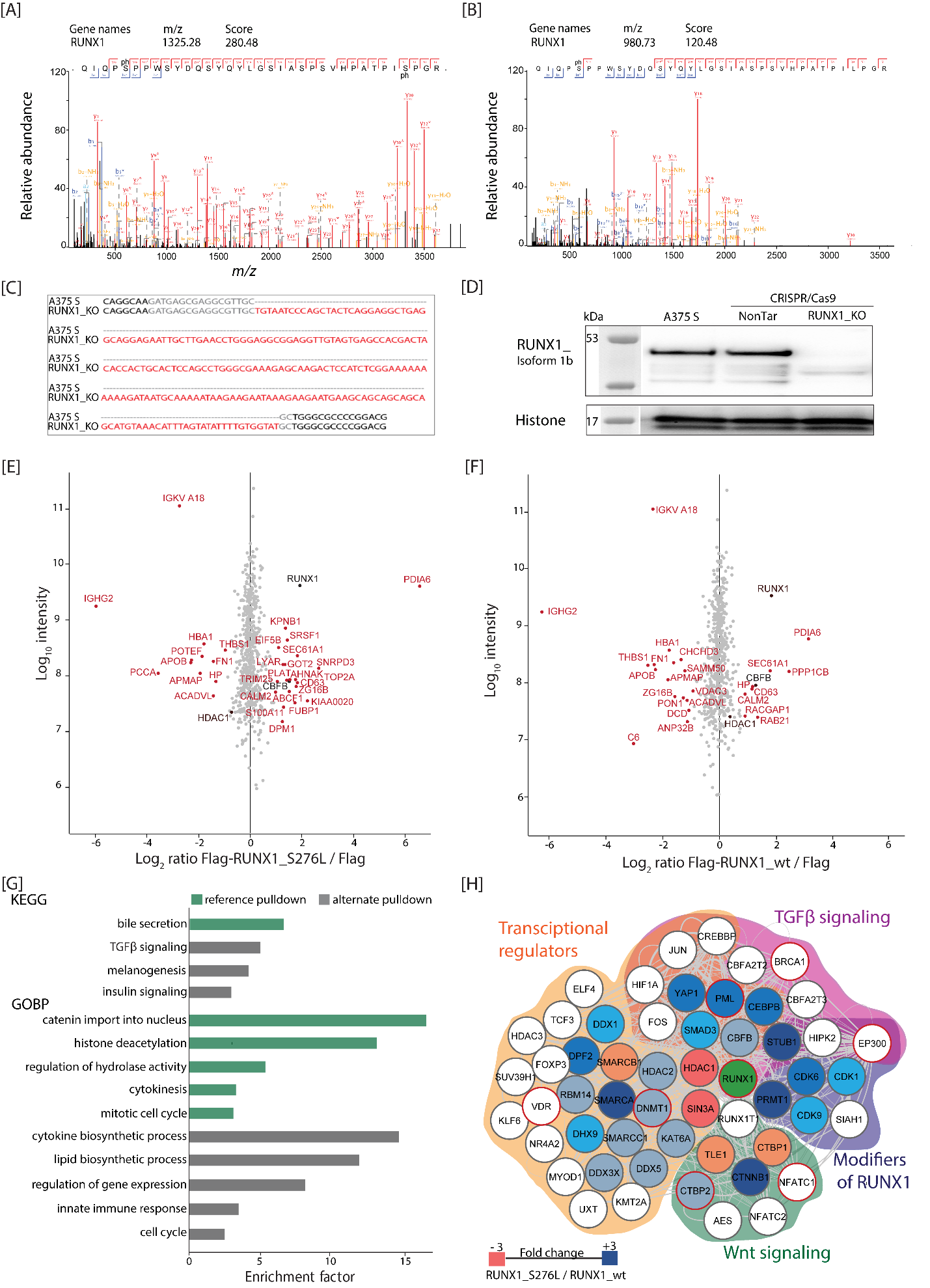

#### Figure S5: Loss of a known phosphorylation site leads to a change in RUNX1 interactome.

[A] and [B] Deconvoluted MS/MS spectrum of phosphorylated reference [A] and alternate [B] peptide of RUNX1 identified by high resolution mass spectrometry. [C] Sanger sequencing result of reference DNA of A375 S and CRISPR/Cas9 genome-edited cell clone RUNX1_KO. [D] Western blot analysis of A375 S, A375 S NonTar and CRISPR/Cas9 genome edited cell clone RUNX1_KO. [E] and [F] Interaction proteomics screen in A375 RUNX1_KO cells stably overexpressing Flag-tagged RUNX1_wt or Flag-tagged RUNX1_S276L. SILAC protein expression (log_2_) of Flag-tagged RUNX1_S276L [E] or Flag-tagged RUNX1_wt [F] relative to the corresponding control cell line (Flag tag only). RUNX1 and its core binding factor CBFB are marked in black. Significantly up and downregulated proteins are highlighted in red. Results represent three replicates per experiment group. [G] Overrepresentation of KEGG pathways and GOBP for reference (Syn_Ser; green) and alternate (Syn_Leu; grey) peptide pulldown. The enrichment factor calculated by Fisher exact test were plotted against the -log_10_ p-value (FDR ≤ 0.1). [H] Protein-protein interaction network for RUNX1 based on BioGRID. Genes identified with nucleotide variant by exome sequencing are circled in red. The node colour correlates with the ratio between RUNX1_S276 and RUNX1_wt. Nodes with white colour are not identified in this study. Overrepresented pathways are coloured as indicated.
